# Practical Model Selection for Prospective Virtual Screening

**DOI:** 10.1101/337956

**Authors:** Shengchao Liu, Moayad Alnammi, Spencer S. Ericksen, Andrew F. Voter, Gene E. Ananiev, James L. Keck, F. Michael Hoffmann, Scott A. Wildman, Anthony Gitter

**Affiliations:** Department of Computer Sciences, University of Wisconsin-Madison; Morgridge Institute for Research; Small Molecule Screening Facility, University of Wisconsin Carbone Cancer Center; Department of Biomolecular Chemistry, University of Wisconsin School of Medicine and Public Health; McArdle Laboratory for Cancer Research, University of Wisconsin-Madison; Department of Biostatistics and Medical Informatics, University of Wisconsin-Madison

**Author notes:** Co-first author (Shengchao Liu, Moayad Alnammi).

## Abstract

Virtual (computational) high-throughput screening provides a strategy for prioritizing compounds for experimental screens, but the choice of virtual screening algorithm depends on the dataset and evaluation strategy. We consider a wide range of ligand-based machine learning and docking-based approaches for virtual screening on two protein-protein interactions, PriA-SSB and RMI-FANCM, and present a strategy for choosing which algorithm is best for prospective compound prioritization. Our workflow identifies a random forest as the best algorithm for these targets over more sophisticated neural network-based models. The top 250 predictions from our selected random forest recover 37 of the 54 active compounds from a library of 22,434 new molecules assayed on PriA-SSB. We show that virtual screening methods that perform well in public datasets and synthetic benchmarks, like multi-task neural networks, may not always translate to prospective screening performance on a specific assay of interest.

## 1 Introduction

Drug discovery is time consuming and expensive. After a specific protein or mechanistic pathway is identified to play an essential role in a disease process, the search begins for a chemical or biological ligand that can perturb the action or abundance of the disease target in order to mitigate the disease phenotype. A standard approach to discover a chemical ligand is to screen thousands to millions of candidate compounds against the target in biochemical or cell-based assays via a process called high-throughput screening (HTS), which produces vast sets of valuable pharmacological data. Even though HTS assays are highly automated, screens of thousands of compounds sample only a small fraction of the millions of commercially-available, drug-like compounds. Cost and time preclude academic laboratories and even pharmaceutical companies from blindly testing the full set of millions of drug-like compounds in HTS assays. Thus, there is a crucial need for an effective virtual screening (VS) process as a preliminary step in prioritizing compounds for HTS assays.

Virtual screening comprises two categories: structure-based^1,2^ and ligand-based methods.^3,4^ Structure-based methods require that the target protein’s molecular structure be known so that the 3D interactions between the target and each chemical compound (binding poses) may be predicted *in silico*. These interactions are given numerical scores, which are then used to rank compounds for potential binding to the target. These methods do not require or typically make use of historical screening data in compound scoring. In contrast, ligand-based methods require no structural information about the target but use data generated from testing molecules in biochemical or functional assays of the target to fit empirical models that relate compound attributes to assay outcomes.

For targets with abundant assay data or where a druggable binding site is not well-defined, such as the targets considered here, ligand-based methods are generally superior to structure-based methods.^5–7^ Confronted with the variety of ligand-based model building methods (e.g., regression models, random forests, support vector machines, etc.),^8^ compound input representations, and performance metrics, how should one proceed? The Merck Molecular Activity Challenge^9^ incited the development of many ligand-based deep learning VS methods,^10-14^ as recently reviewed.^15,16^ These methods are often assessed with cross-validation on existing HTS data, but there is presently little experimental evidence on the best option for prioritizing new compounds given a fixed screening budget.

We critically evaluated a collection of VS algorithms that include both structure-based and ligand-based methods, with a focus on the subset of quantitative structure-activity relationship ligand-based methods that use machine-learning to predict active compounds for a target based on initial screening data. We present a VS workflow that first uses available HTS training data to systematically prune the specific versions of the algorithms and calculate their cross-validation performance on a variety of evaluation metrics. Based on the cross-validation results and analysis of the various evaluation metrics, we selected a single virtual screening algorithm. The selected method, a random forest model, was the best option for prioritizing a small number of compounds from a new library, as verified by experimental screening. These model selection and evaluation strategies can guide VS practitioners to select the best model for their target even as the landscape of available VS algorithms continues to evolve.

## 2 Methods

### 2.1 Datasets

Our case studies were on new and recently generated datasets^17,18^ for the targets PriA-SSB and RMI-FANCM. The PriA-SSB interaction is important in bacterial DNA replication and is a potential target for antibiotics.^19^ The RMI-FANCM interaction is involved in DNA repair that is induced in human cancer cells to confer chemoresistance to cytotoxic DNA-cross-linking agents, making it an attractive drug target.^20^ We previously screened these targets with a library of compounds obtained from Life Chemicals Inc. (LC) in different assay formats. In addition, we screened new LC compounds on the PriA-SSB target to evaluate our VS models. The four datasets derived from these screens are described below and summarized in Table 1.

**Table 1:** Summary statistics for the four binary datasets.

| Stage | Dataset | % inhibition threshold | # actives | # inactives |
| --- | --- | --- | --- | --- |
| Cross-validation | **PriA-SSB AS** | $\geq 35\%$ | 79 | 72,344 |
| | **PriA-SSB FP** | $\geq 30\%$ | 24 | 72,399 |
| | **RMI-FANCM FP** | $\geq \text{mean} + 2 \text{ SD}$ | 230 | 49,566 |
| Prospective | **PriA-SSB prospective** | $\geq 35\%$ | 54 | 22,380 |

#### PriA-SSB AlphaScreen

PriA-SSB was initially screened using an AlphaScreen (AS) assay in 1536-well format^18^ on 72,423 LC compounds at a single concentration (33.3 µM), with data reported as % inhibition compared to controls. We refer to these continuous values as “PriA-SSB %inhibition”. Those compounds that tested above an activity threshold (*≥* 35% inhibition) and passed PAINS chemical structural filters^21,22^ were retested in the same AS assay. Although PAINS filters are not a technical necessity of any VS method, it is now a common requirement for publication of HTS and medicinal chemistry projects. In this study, we apply PAINS filters for the internal consistency of the data sets. Compounds that were confirmed in the AS retest screen (*≥* 35% inhibition) were marked as actives, creating the binary dataset **PriA-SSB AS**.

##### PriA-SSB fluorescence polarization

Compounds that had PriA-SSB % inhibition *≥* 35% and passed the PAINS filters were also tested in a fluorescence polarization (FP) assay as a secondary screen. Those compounds with FP inhibition *≥* 30% were labeled as actives, creating the binary dataset **PriA-SSB FP**, with all other compounds in the screening set labeled inactive.

##### RMI-FANCM fluorescence polarization

The RMI-FANCM interaction was initially screened with a subset of 49,796 compounds from the same LC library as PriA-SSB.^17^ This FP assay was run at a single compound concentration (32 µM). We refer to these continuous values as “RMI-FANCM % inhibition”. Those compounds that demonstrated activity *≥* 2 standard deviations (SD) above the assay mean and passed PAINS filters were marked as actives in the binary dataset **RMI-FANCM FP**.

##### PriA-SSB prospective

For prospective testing, we experimentally screened an additional 22,434 compounds after the VS methods predicted their activity. We removed compounds that were already included in the 72,423 LC compounds in the **PriA-SSB AS** dataset to ensure there was no overlap between the prospective screen compounds and those used to train VS models. As with the initial library, the PriA-SSB AS assay was used in the same 1536-well format at a single concentration (33.3 µM) to test the additional 22,434 LC compounds. Actives were defined with the same criteria used for the binary dataset **PriA-SSB AS**. Compounds with at least 35% inhibition that passed the PAINS filters were retested with the AS assay. Those with at least 35% inhibition in the AS retest were labeled as actives, creating the binary dataset **PriA-SSB prospective**.

Because secondary screens and structural filters were used to define the active compounds, there was no single primary screen % inhibition threshold that separated the actives from the inactives. Some compounds exhibiting high % inhibition values were labeled as inactive because they did not satisfy the structural requirements or were not active in the secondary screen.

##### PubChem BioAssay

To help learn a better chemical representation with multi-task neural networks, we considered other screening contexts from which to transfer useful knowledge. We used a subset of 128 assays (AIDs) from the **PubChem BioAssay (PCBA)**^23^ repository. This dataset was used in previous work on multi-task neural networks.^14^ This subset contained assays for which the assays were developed to probe a specific protein target and dose-response measurements were obtained for each compound (see Appendix A for other assay query filters). Potency and curve quality are factored into a PubChem Activity Score. Regardless of assay, compounds with a PubChem Activity Score of 40 or greater (range 0-100) were assigned a PubChem Bioactivity outcome (label) of “Active”. Compounds with PubChem Activity Scores 1-39 were labeled “Inconclusive,” and those with 0 were labeled “Inactive” (Appendices B and C).

### 2.2 Compound Features

Ligand-based virtual screening methods require each chemical compound to be represented in a particular format as input to the model. We adopted two common representations. All of the ligand-based algorithms except the Long Short-Term Memory (LSTM) neural network use 1024-bit Morgan fingerprints^24^ with radius 2 generated with RDKit version 2016.03.4.^25^ These circular fingerprints are similar to ECFP4 fingerprints,^26^ though with a slightly different implementation. For LSTM networks, we used the Simplified Molecular Input Line Entry System (SMILES) representation,^27^ where the characters were treated as sequential features.

### 2.3 Virtual Screening Models

We selected a variety of existing virtual screening approaches for our benchmarks and prospective predictions. These included ligand-based supervised machine learning approaches, structure-based docking, and a chemical similarity baseline. Table 2 summarizes the types of training data used by each algorithm.

**Table 2:** A summary of the virtual screening methods and which labels each model used during training. The docking and consensus docking models do not train on the PriA-SSB or RMI-FANCM datasets.

| Model | Continuous % inhibition | Binary label | PCBA binary labels |
| --- | --- | --- | --- |
| Dock |  |  |  |
| CD |  |  |  |
| STNN-C |  | ✓ |  |
| STNN-R | ✓ |  |  |
| MTNN-C |  | ✓ | ✓ |
| LSTM |  | ✓ |  |
| RF |  | ✓ |  |
| IRV |  | ✓ |  |
| Similarity baseline |  | ✓ |  |

#### 2.3.1 Ligand-Based Neural Networks

Deep learning is a machine learning approach that encompasses neural network models with multiple hidden layer architectures and the techniques for training these models. It represents the state of the art for many predictive tasks, which has generated extensive interest in deep learning for biomedical research, including virtual screening.^15,16^ We evaluated multiple types of established neural network architectures for virtual screening.

##### Single-task Neural Network (STNN)

A single-task neural network (Figure 1(a)) makes a single prediction for a single target (also referred to as a task). We trained a separate model for each of the **PriA-SSB AS**, **PriA-SSB FP**, and **RMI-FANCM FP** datasets, taking each compound’s Morgan fingerprint as the input features. We trained the neural networks using Keras^28^ with the Theano backend.^29^ The single-task neural networks were trained on each task to predict either the binary activity label in the classification setting (STNN-C) or the continuous % inhibition in the regression setting (STNN-R). These neural networks used two hidden layers with 2000 hidden units each, Adam optimization,^30^ 0.25 dropout rate, and other hyperparameters described in Tables S2 and S3.

**Figure 1:**
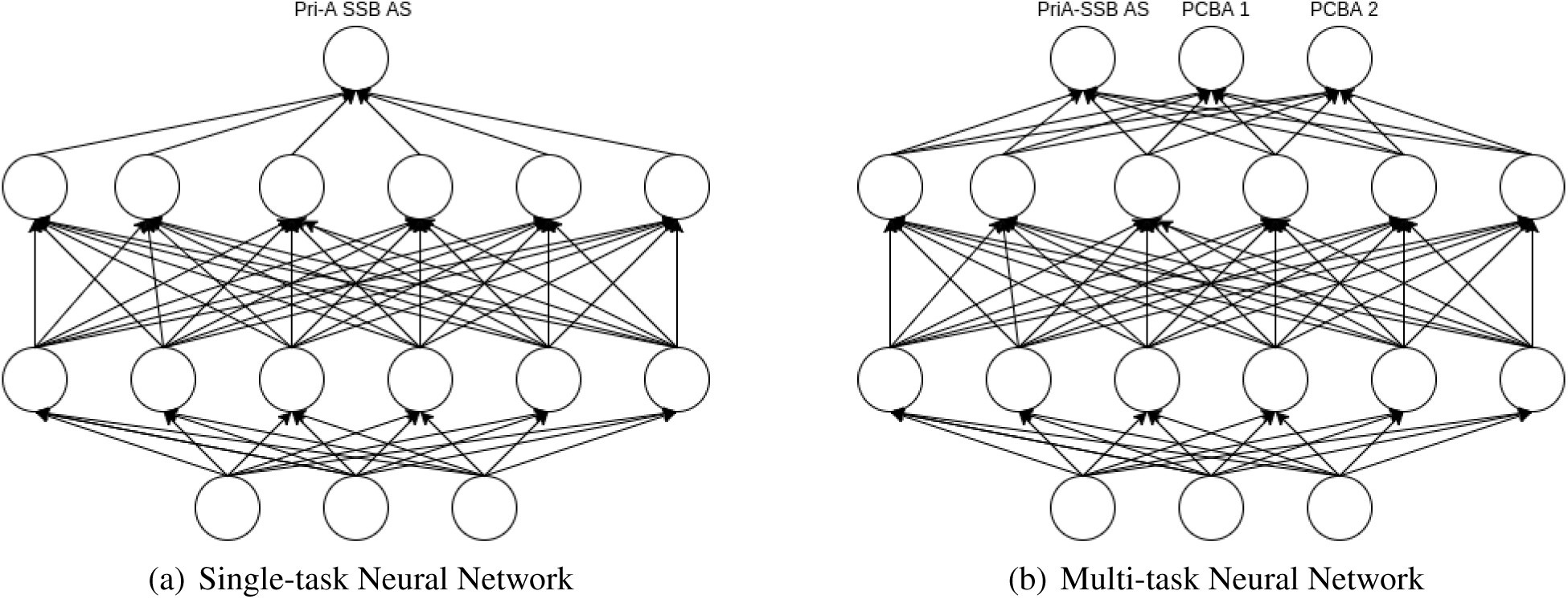
Neural network structures. The neural networks map the input features (e.g. fingerprints) in the input (bottom) layer to intermediate chemical representations in the hidden (middle) layers and finally to the output (top) layer, which makes either continuous or binary predictions. Figure 1(a) has only one unit in the output layer. Figure 1(b) has multiple units in the output layer representing different targets, one for our new target of interest and the others for PCBA targets.

##### Multi-task Neural Network (MTNN)

Multi-task neural networks make different predictions for multiple targets or tasks but share knowledge by training the first few hidden layers together. Each of our multi-task neural networks included one target task (**PriA-SSB AS**, **PriA-SSB FP**, or **RMI-FANCM FP**) and 128 tasks from PCBA. We only trained multi-task neural networks in the classification setting (MTNN-C). The MTNN-C models used two hidden layers with 2000 hidden units each, Adam optimization, 0.25 dropout rate, and other hyperparameters described in Table S2.

##### Single-task Atom-level LSTM (LSTM)

The LSTM is one of most prevalent recurrent neural network models,^31^ which has been applied previously in virtual screening.^32^ An LSTM assumes there exists a sequential pattern in the input string. We used a one hot encoding of the SMILES strings as input for the LSTM model. In a one hot encoding, each character in a SMILES string is replaced by a binary vector. The binary vector has one bit for each possible unique character in all SMILES strings. At each position in a SMILES string, the bit corresponding to the character at that position is set to 1, and all other bits are set to 0. We trained the LSTM model to predict the binary activity labels. The LSTM models used one or two hidden layers with 10 to 100 hidden units each, Adam optimization, 0.2 or 0.5 dropout rate, and other hyperparameters described in Table S4. The compounds in the cross-validation stage used SMILES generated by OpenEye Babel version 3.3. The compounds in the prospective screen were processed separately and used SMILES from RDKit version 2016.03.4.^25^

##### Influence Relevance Voter (IRV)

IRV^33,34^ is a hybrid between *k*-nearest neighbors and neural networks. Each compound’s output value is a non-linear combination of the similarity scores from its most closely related compounds in the training dataset. We used Morgan fingerprints as the input and trained separate IRV models for each dataset. The IRV models used 5 to 80 neighbors and other hyperparameters described in Table S5.

#### 2.3.2 Other Ligand-Based Models

##### Random Forest (RF)

Random forests are ensembles of decision trees that are often used as a baseline in virtual screening benchmarks.^35,36^ We used scikit-learn^37^ to train a random forest classifier for each binary label with Morgan fingerprints as features. The RF models used 4,000 to 16,000 estimators, 1 to 1000 minimum samples at a leaf node, a bound on the maximum number of features, and other hyperparameters described in Table S6.

#### 2.3.3 Protein-Ligand Docking

##### Target Preparation

Our structure-based VS approach involved the docking-based ranking of the LC library to the holo-form of PriA using the crystal structure (PDB: 4NL8),^48^ in which it is bound to a C-terminal segment of an SSB protein. A missing loop in this structure was added from the the apo-form (PBD: 4NL4), though this is not near the SSB binding site. The docking search space was limited to 8Å from the coordinates of the co-crystallized SSB C-terminal tripeptide.

For RMI-FANCM, the RMI protein was built from both the A and B chains from the structure (PDB: 4DAY).^39^ The docking search space was defined by the central 5 residues of the MM2 peptide (PDB: 4DAY chain C), Val-Thr-Phe-Asp-Leu, also with an 8 Å bounding box.

##### Compound Preparation

LC library compounds were assigned 3D coordinates and Merck Molecular Force Field partial charges using OpenEye OMEGA and Molcharge.^40^ Compounds in the LC library with ambiguous stereochemistry were enumerated in all possibilities, and the best resulting docking score was retained for each.

##### Docking (Dock) and Consensus Docking (CD)

We ran eight different docking programs and generated nine docking scores as a broad comparison to the ligand-based methods under consideration. The docking programs and names we use for their scores are AutoDock version 4.2.641 (Dock_ad4), Dock version 6.742 (Dock_dock6), FRED version 3.0.143 (Dock_fred), HYBRID version 3.0.1^43^ (Dock_hybrid), PLANTS version 1.2^44^ (Dock_plants), rDock version 2013.1^45^ (Dock_rdocktot and Dock_rdockint), Smina version 1.1.2^46^ (Dock_smina), and Surflex-Dock version 3.040^47^ (Dock_surflex). In addition, we calculated consensus docking scores using three traditional approaches (CD_mean, CD_median, and CD_max) and two versions of the Boosting Consensus Score (CD_efr1_opt and CD_rocauc_opt).^48^ The consensus docking methods were developed without any knowledge of the PriA-SSB or RMI-FANCM assay data. Compounds with missing scores due to preparation or docking failures were not considered during evaluation.

#### 2.3.4 Chemical Similarity Baseline

In the prospective screening stage, we introduced a simple baseline compound ranking method against which to compare the more sophisticated VS methods. The active compounds from **PriA-SSB AS** were used as seeds for similarity searching through all 22,434 compounds in the **PriA-SSB prospective** dataset. The prospective set compounds were ranked by their maximum Tanimoto similarity to any of the **PriA-SSB AS** actives with MayaChemTools^49^ using Morgan fingerprints from RDKit version 2013.09.1.

In addition, all compounds were clustered by two separate approaches to describe chemical series. Chemical similarity-based hierarchical clusters on Morgan fingerprints using Ward’s clustering are described as SIM. Maximum common substructure clusters, used to group molecules with similar scaffolds, are described as MCS. JKlustor was used for both types of clustering (JChem version 17.26.0, ChemAxon).

### 2.4 Evaluation Metrics

Given our goal of developing VS methods that enable very small, cost-effective, productive screens, we considered how metrics weight early active retrieval. All of the VS algorithms produce a ranked list of compounds, where compounds are ordered by the probability of being active, the continuous predicted % inhibition, the docking score, or a comparable output value. For a ranked list of copounds, we can threshold the ranked list and consider all compounds above the threshold as positive (active) predictions and those below the threshold as negative (inactive). Classification models output class probabilities. Regression models, docking, and the similarity baseline outputs different types of continuous scores. Thresholding on the compound rank is equivalent to thresholding on the class probability or continuous score because for each rank there is a corresponding probability or score. By comparing those predictions to the experimentally-observed activity, we can compute true positive (TP), false positive (FP), true negative (TN), and false negative (FN) predictions for the ranked list at that threshold. We explored several options for summarizing how well each algorithm ranks the known active compounds. Because most of the compounds have only single-replicate measurements of % inhibition, we focus on evaluating active versus inactive compounds instead of correlation with the % inhibition.

The area under the receiver operating characteristic curve (AUC[ROC]) has been recommended for virtual screening because it is robust, interpretable, and does not depend on user-defined parameters.50 The ROC curve plots the relationship between true positive rate (TPR, also known as sensitivity or recall) and false positive rate (FPR, equivalent to 1 - specificity), which are defined in Equation 1. As the FPR goes to 100%, all ROC curves converge, whereas early active retrieval (a more meaningful characteristic of VS performance) can be assessed in the low FPR region of the ROC curve, which exhibits greater variability across VS methods. Thus, we also considered the Boltzmann-enhanced discrimination of receiver operating characteristic (BEDROC).^51^ It emphasizes the early part of the ROC curve through a scaling function *α*, which we set to 10 for our purposes of early enrichment up to 20%. We used the BEDROC implementation from the CROC Python package.^52^

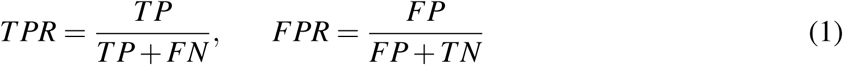

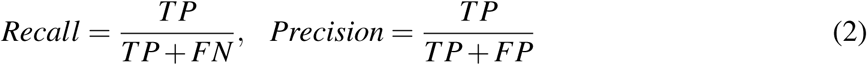

Area under the precision-recall curve (AUC[PR]) is another common metric (Equation 2). AUC[PR] has an advantage over AUC[ROC] for summarizing classifier performance when the class labels are highly-skewed, as in virtual screening where there are few active compounds in a typical library. AUC[PR] evaluates a classifier’s ability to retrieve actives (recall) and which of the predicted actives are correctly classified (precision) as we vary the prediction threshold. We used the PRROC R package’s “auc.integral”^53^ to compute AUC[PR].

Another VS metric is enrichment factor (EF), which is the ratio between the number of actives found in a prioritized subset of compounds versus the expected number of actives in a random subset of same size. In other words, it assesses how much better the VS method performs over random compound selection. Let *R* ∈ [0%, 100%] be a pre-defined fraction of the compounds from the total library of compounds screened.

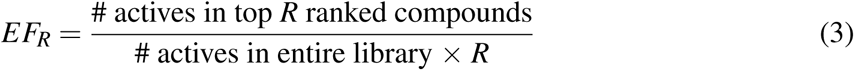

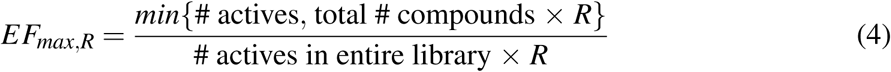

*EF*_*max, R*_ represents the maximum enrichment factor possible at *R*. Difficulty arises when interpreting EF scores because they vary with the dataset and threshold *R*. We defined the normalized enrichment factor (NEF) as:

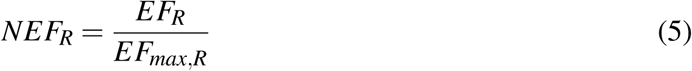

Because *NEF*_*R*_ ∈ [0, 1], it is easier to compare performance across datasets and thresholds. 1.0 is the perfect NEF. Furthermore, we can create an NEF curve as *NEF*_*R*_ versus *R ∈* [0%, 100%] and compute the area under that curve to obtain AUC[NEF] ∈ [0, 1]. However, most models tend to exhibit similar late enrichment behavior. We are typically interested in early enrichment behavior so we computed AUC[NEF] using *R* ∈ [0%, 20%].

Finally, we introduce the metric *n*_*hits*_, which is simply the number of actives found in a selected number of tested compounds (e.g. how many hits or actives were found in 250 tested compounds). This metric represents the typical desired utility of a screening process: retrieve as many actives as possible in the selected number of tests (denoted as *n*_*tests*_). We compare *n*_*hits*_ at various *n*_*hits*_ to the different evaluation metrics to identify which metrics best mimic the *n*_*hits*_ utility.

### 2.5 Pipeline

Our virtual screening workflow contains three stages:

1. Tune hyperparameters in order to prune the model search space.
2. Train, evaluate, and compare models with cross-validation to select the best models.
3. Assess the best models’ ability to prospectively identify active compounds in a new set.

In contrast to most other virtual screening studies, the experimental screen was not conducted until after all models were trained and evaluated in the cross-validation stage (Figure 2). For the first two stages, we first split the **PriA-SSB AS**, **PriA-SSB FP**, and **RMI-FANCM FP** datasets into five stratified folds as described in Appendix B.

**Figure 2:**
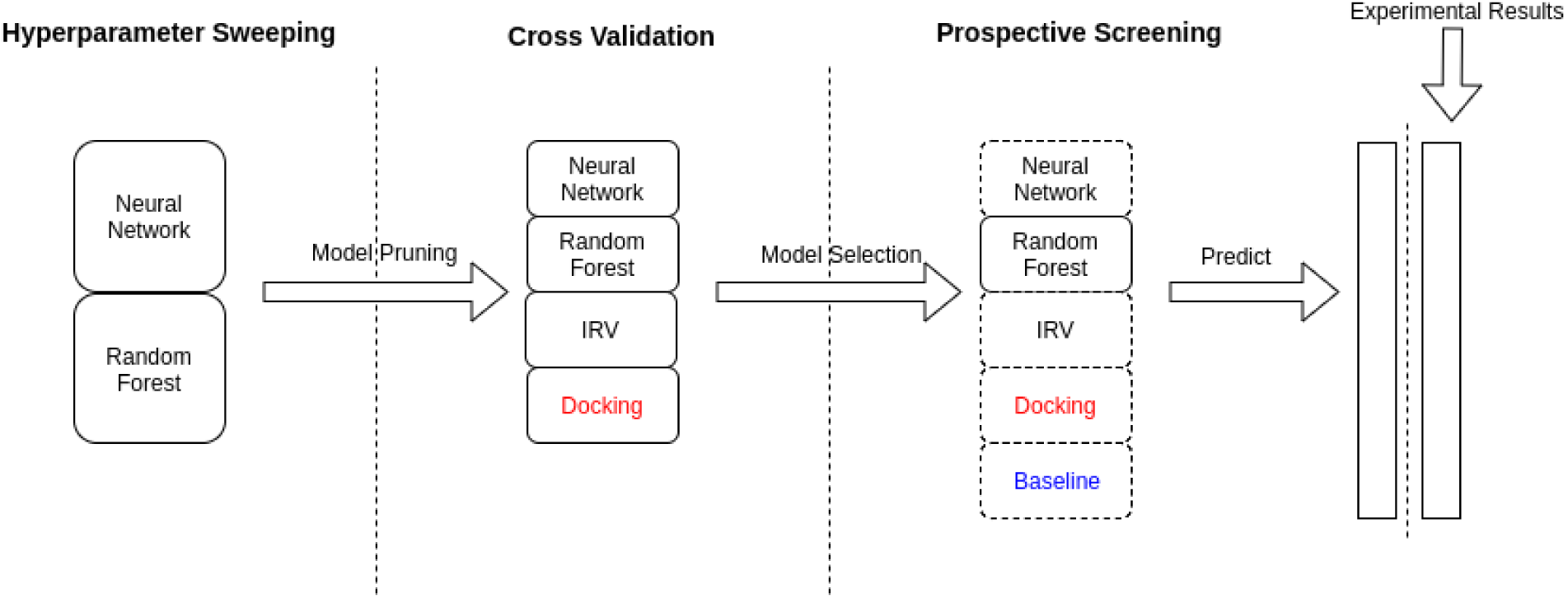
Initially, 258 neural network and random forest models are evaluated to eliminate poorly performing hyperparameter combinations. The models with the best hyperparameters advance to cross-validation along with IRV and docking-based methods, for a total of 35 models. Cross-validation identifies a random forest as the best overall model. The VS methods and similarity baseline then predict active compounds in the **PriA-SSB prospective** dataset. After the predictions are finalized, we experimentally screen the compounds to evaluate the predictions. Black text denotes ligand-based machine learning models, red text denotes docking-based models, and blue text is the similarity baseline.

#### 2.5.1 Hyperparameter Sweeping Stage

Hyperparameters are model configurations or settings that are set by an expert as opposed to the weights or parameters that are learned or fit during model training. For most of the ligand-based machine learning models, the hyperparameter space was too large for exhaustive searches using the full dataset. Therefore, we applied a grid search on a pre-defined set of hyperparameters in a smaller dataset and pruned those that performed poorly. We performed a single iteration of training on the first four folds of **PriA-SSB AS** to avoid over-fitting. The hyperparameters considered are listed in Appendix D.

#### 2.5.2 Cross-Validation Stage

To identify which VS algorithms are likely to have the best performance in a prospective screen, we applied a traditional cross-validation training strategy on datasets **PriA-SSB AS**, **PriA-SSB FP**, and **RMI-FANCM FP** after reducing the hyperparameter combinations to consider. Selecting the best model is non-trivial. Ideally, the best model would have dominant performance on all evaluation metrics, but this is rarely observed with existing models. Each evaluation metric prioritizes different performance characteristics. Our cross-validation results illustrate which models consistently perform well over different metrics, the correspondence of metrics relative to a desired utility (*nhits*), and how to choose models and evaluation metrics in order to successfully identify active compounds in a prospective screen.

Cross-validation is commonly used to avoid over-fitting when there are few training samples. We split the training data into 5 folds, 4 folds for training and 1 for testing. Models like RF and IRV that do not require a hold-out dataset for early stopping used 4 folds for training. The neural networks perform early-stopping based on a hold-out set, so we iteratively selected 1 of the 4 training data folds for this purpose. This led to a nested cross-validation with 5 *×* 4 = 20 trained neural networks.

#### 2.5.3 Prospective Screening Stage

Our prospective screen used a library of 22,434 new LC compounds that were not present in the **PriA-SSB AS** training set. We used each VS model to prioritize 250 of these compounds that are most likely to be active. This emulates virtual screening on much larger compound libraries, in which only a small fraction of all computationally scored compounds can be tested experimentally. When models assigned the same score to multiple compounds, we broke ties arbitrarily to obtain exactly 250 compounds.

After finalizing the models’ predictions, we screened all 22,434 compounds in the wet lab and assigned actives based on a 35% inhibition threshold and structural filters (the **PriA-SSB prospective** dataset). Finally, we evaluated how many of the experimental actives each VS method identified in its top 250 predictions, the number of distinct chemical clusters recovered, and the number of active compounds that were not in the top 250 predictions from any of the VS algorithms. The prospective screen allowed us to assess how well the cross-validation results generalized to new compounds and further verified our conclusions from the retrospective cross-validation tests.

### 2.6 Data and Software Availability

Code implementing our ligand-based virtual screening algorithms is available at https://github.com/gitter-lab/pria_lifechem. This GitHub repository also contains additional Jupyter notebooks to reproduce the visualizations and analyses. Version 0.1.0 of our software is archived on Zenodo (doi:10.5281/zenodo.1257674). Our new PriA-SSB HTS data are available on PubChem (PubChem AID:1272365) along with the existing RMI-FANCM HTS data (PubChem AID:1159607). A formatted version of this dataset for training virtual screening algorithms is available on Zenodo (doi:10.5281/zenodo.1257462).

## 3 Results

### 3.1 Cross-Validation Results

In the cross-validation stage we assessed 35 models: 8 neural networks (STNN-C (Table S7), STNN-R (Table S8), MTNN-C (Table S9), and LSTM (Table S10)), 5 IRV (Table S11), 8 RF (Table S12), and 14 from docking (Dock) or consensus docking (CD). When there are multiple versions of a model that use different hyperparameters, we distinguish them with alphabetic suffixes such as “_a” and “_b”. Tables S7-S12 describe the hyperparameters associated with these suffixes. We highlight the **PriA-SSB AS** dataset as a representative example, but the VS workflow is applicable for all tasks.

#### 3.1.1 Comparing Virtual Screening Algorithms

We tested all 35 models on three datasets, and the results for four evaluation metrics on the **PriA-SSB AS** dataset are shown in Figure 3. Appendix F contains the results for **PriA-SSB FP** and **RMI-FANCM FP**. The **PriA-SSB AS** performance using AUC[ROC] was comparable for many models. All models except LSTM, some IRV models, and docking were above 0.8 AUC[ROC]. Random forest was the best model, especially for the metrics that prioritize early enrichment.

**Figure 3:**
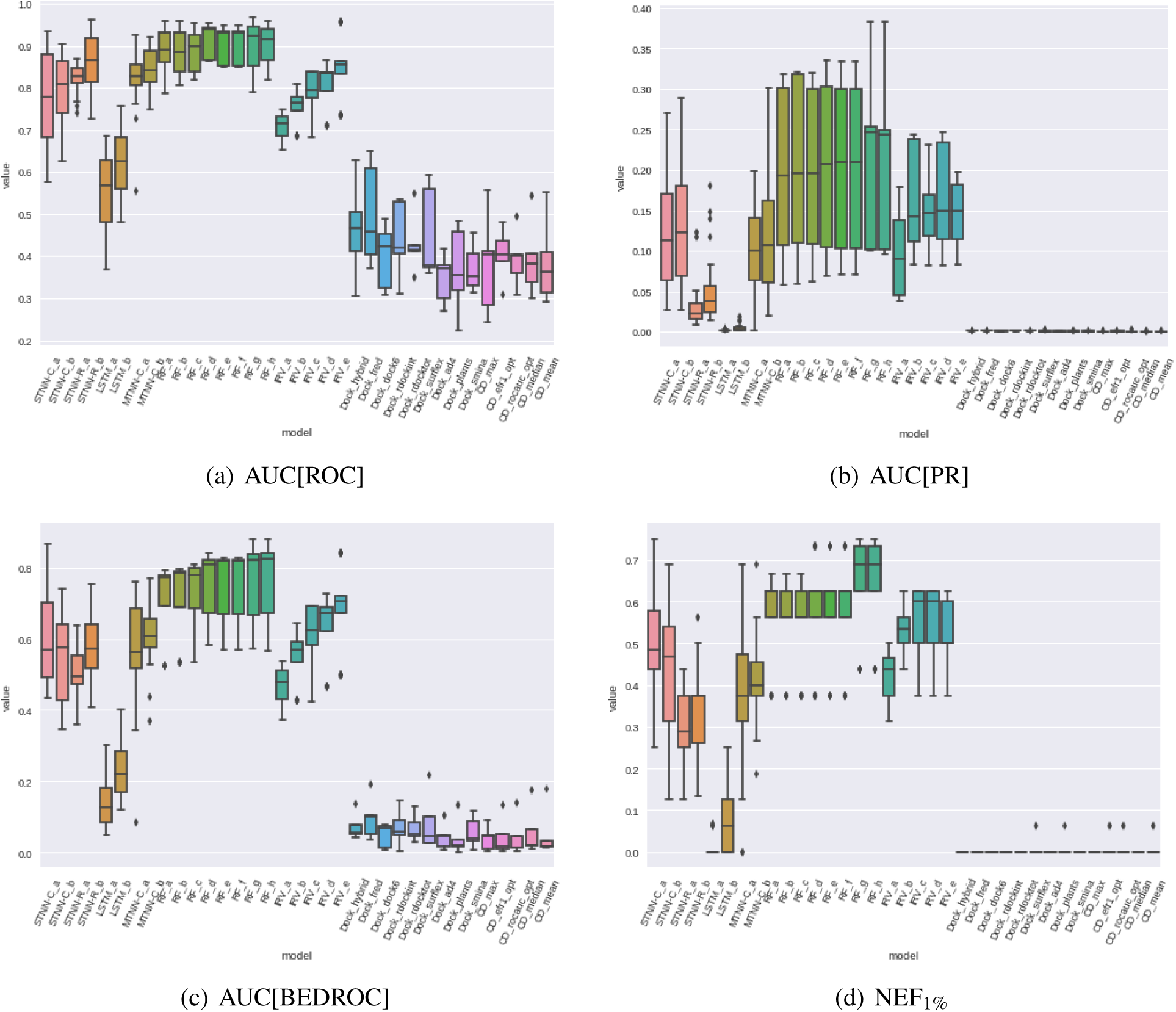
The evaluation metric distributions on **PriA-SSB AS** on all models over the cross-validation folds. The metrics are: (a) AUC[ROC], (b) AUC[PR], (c) AUC[BEDROC], and (d) NEF^1%^ as described in section 2.4.

Random forest was again the best overall method for the **RMI-FANCM FP** dataset (Appendix F). On the **PriA-SSB FP** dataset, STNN-R achieved the highest scores over the majority of the metrics (Appendix F). The other types of VS models were effectively tied for most metrics (Appendix G).

#### 3.1.2 Evaluation Metrics

Given a fixed evaluation metric, we could compare two models with a t-test to assess if one statistically outperforms the other. However, we needed to make such comparisons repeatedly between each pair of models and required a statistical test that accounts for multiple hypothesis testing. Due to unequal variances and sample sizes (Figure 3), we used Dunnett’s modified Tukey-Kramer test (DTK)^54,55^ for pairwise comparison to assess whether the mean metric scores of two models were significantly different. Using DTK results for *each metric*, we scored each model based on how many times it attained a statistically significantly better result than other models (Appendix G). For most metric-target pairs, many models have the same rank because DTK does not report a significant difference.

In a prospective screen, our goal is to maximize the number of active compounds identified by a VS algorithm given a fixed budget (number of predictions). We wanted to determine which of the VS evaluation metrics best aligns with *n*_*hits*_. Thus, we compared the model ranking induced by each metric with the model ranking induced by *n*_*hits*_ at varying number of tests.

To score the evaluation metrics, we used Spearman’s rank correlation coefficient based on the model rankings induced by the metric of concern versus *n*_*hits*_ at a specific *n*_*tests*_. We then ranked the metrics based on their correlation with *n*_*hits*_. The correlation coefficients and metric rankings can be found in Appendix H. The metric ranking varies depending on *n*_*tests*_ and the target. Some metrics overtake one another as we increase or decrease *n*_*tests*_ For **PriA-SSB AS**, NEF_*R*_ consistently placed in the top ranking correlations when *R* coincided with *n*_*tests*_. This is evident when we focus on a single metric and see the top ranking metrics for *n*_*tests*_ ∈ [100, 250, 500, 1000, 2500]. Only for a large enough *n*_*tests*_ do metrics like AUC[ROC] that evaluate the complete ranked list become comparable. This suggests that if we know *a priori* how many new compounds we can afford to screen, then NEF_*R*_ at a suitable *R* is a viable metric for choosing a VS algorithm during cross-validation in the hopes of maximizing *n*_*hits*_.

#### 3.1.3 Selecting the Best Model

Based on these results, we selected the VS screening models that are most likely to generalize to new compounds and identify actives in our experimental screen of 22,434 new compounds. We focus on PriA-SSB for the prospective screen using models trained on **PriA-SSB AS** because the assay was more readily available for us to generate data for the new compounds.

Table 3 compares model selection based on evaluation metric means alone versus the DTK+Mean approach for multiple evaluation metrics on the three tasks. The complete model rankings for means only and DTK+Mean can be found in Appendix I. DTK+Mean ranks models by statistical significance and uses the mean value only for tie-breaking. Both strategies selected the same models for a fixed evaluation metric, except for AUC[PR] on all three tasks and NEF_1%_ for **RMI-FANCM FP** (Table 3). This is mainly due to DTK not detecting statistically significant differences among the models’ evaluation scores, so tie-breaking by means selected the same models as ranking by means. Recall that **PriA-SSB FP** has fewer actives than **PriA-SSB AS** and **RMI-FANCM FP** (Table 1). Similar RF and STNN-C models were selected for **PriA-SSB AS** and **RMI-FANCM FP**. However, **PriA-SSB FP** identified STNN-R models exclusively.

**Table 3:** Top-ranked models by means versus DTK+Mean on the three tasks. Evaluation metric means were computed over all cross-validation folds. The prospective screening was only performed on PriA-SSB. Model names are mapped to their hyperparameter values in Appendix E.

| Metric | Best by Mean Model |  |  | Best by DTK+Mean Model |  |  |
| --- | --- | --- | --- | --- | --- | --- |
|  | PriA-SSB AS | PriA-SSB FP | RMI-FANCM FP | PriA-SSB AS | PriA-SSB FP | RMI-FANCM FP |
| AUC[ROC] | RF_d | STNN-R_a | RF_h | RF_d | STNN-R_a | RF_h |
| AUC[BEDROC] | RF_h | STNN-R_b | RF_h | RF_h | STNN-R_b | RF_h |
| AUC[PR] | RF_g | STNN-R_a | RF_h | STNN-C_b | STNN-R_b | STNN-C_b |
| AUC[NEF] | RF_h | STNN-R_b | RF_h | RF_h | STNN-R_b | RF_h |
| NEF<sub>1%</sub> | RF_h | STNN-R_b | RF_h | RF_h | STNN-R_b | RF_h |

In our prospective screen, each model prioritizes 250 top-ranked compounds, approximately 1% of the new LC library. In this setting where each model has a fixed budget for the predicted compounds, NEF_*R*_ is a suitable metric. Therefore, we used NEF_1%_ with DTK+Means to choose the best models from each class. The best-in-class models were RandomForest_h, SingleClassification_a, SingleRegression_b, MultiClassification_b, LSTM_b, IRV_d, and ConsensusDocking_efr1_opt, with RandomForest_h being the strongest model overall (Appendix I).

### 3.2 Prospective Screening Results

After selecting the best model from each class based on cross-validation and the NEF_1%_ metric, we retrained the models on all 72,423 LC compounds to predict PriA-SSB inhibition using the same types of data shown in Table 2. This provided a single version of each model instead of one for each cross-validation fold. All models then ranked 22,434 new LC compounds that were provided without activity labels. We selected the top 250 ranked new compounds from each model. Then, we experimentally screened all 22,434 new compounds to assess PriA-SSB % inhibition and defined actives based on a 35% inhibition threshold and PAINS filters. The new binary dataset **PriA-SSB prospective** contained 54 actives.

Table 4 presents how many of the 54 actives were identified by each best-in-class virtual screening method and the chemical structure similarity baseline. For comparison, randomly selecting 250 compounds from the **PriA-SSB prospective** dataset is expected to identify less than one active based on the overall hit rate. Appendices J and K show the VS models’ **PriA-SSB prospective** performance for the other evaluation metrics.

**Table 4:**
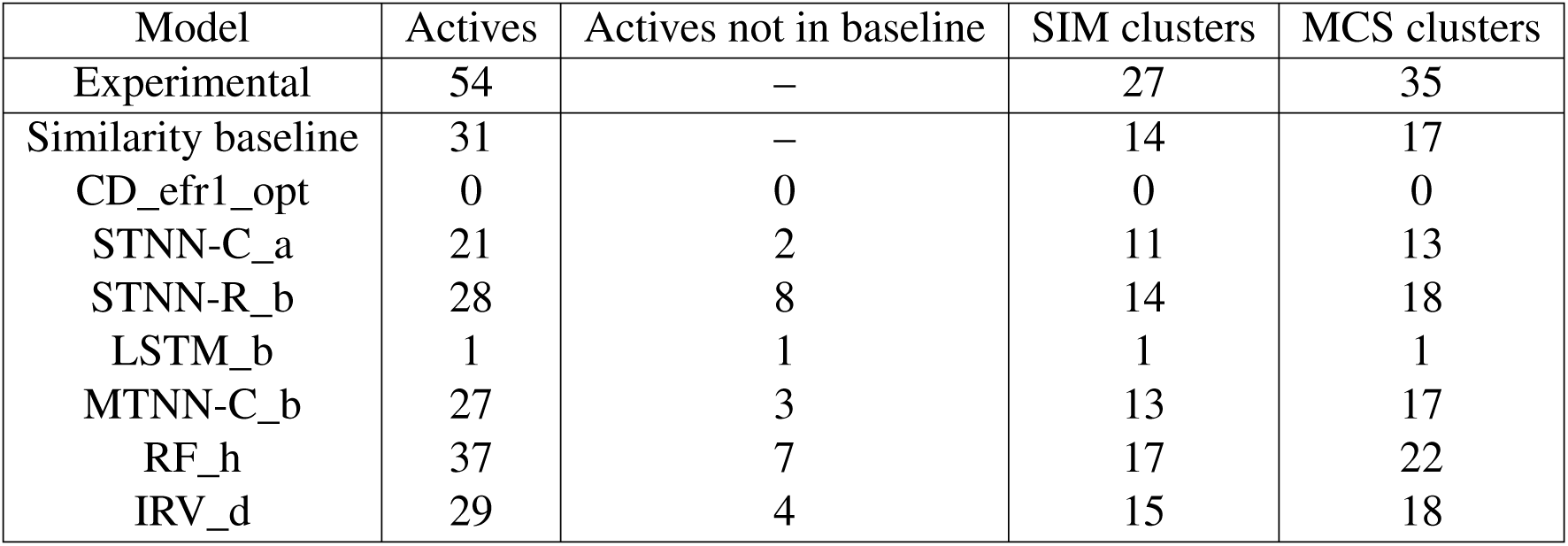
The number of active compounds in the top 250 predictions from the seven selected models and the chemical similarity baseline compared to the number of experimentally-identified actives. These selected models are the best in each algorithm category from cross-validation. The last two columns correspond to the number of distinct chemical clusters from similarity or maximum common substructure clustering that are represented among the 54 actives. Prospective performance for all VS models is in Appendix L.

Table 4 also lists the number of distinct chemical clusters identified by each method, with the goal of identifying as many diverse active compounds as possible. The 22,434 compounds form 124 SIM and 714 MCS clusters or chemical series. Of these, the 54 experimental actives represent 27 SIM and 35 MCS clusters. Commonly, virtual screening is followed by a medicinal chemistry effort that would be expected to identify other members of these clusters.

In general, the number of distinct chemical clusters captured in the top 250 predictions is correlated with the number of actives (Table 4), meaning that the methods selected structurally diverse hits. The similarity baseline identified compounds from roughly half of the SIM or MCS clusters. With the exceptions of docking and LSTM_b, each of the methods in Table 4 found at least one cluster not present in the baseline. The machine learning techniques are not limited to finding only the chemotypes that are present in the training set (Appendix M).

The ligand-based VS methods recovered many of the same actives as the chemical similarity baseline, but they also found actives that were missed by the baseline (Figure 4). There was a group of eleven active compounds that were identified by most ligand-based methods, including the baseline model. The compounds identified are not the most potent, either within their cluster or overall. Nor do any of the methods exhibit any correspondence between the number of compounds identified from a cluster and their potency.

**Figure 4:**
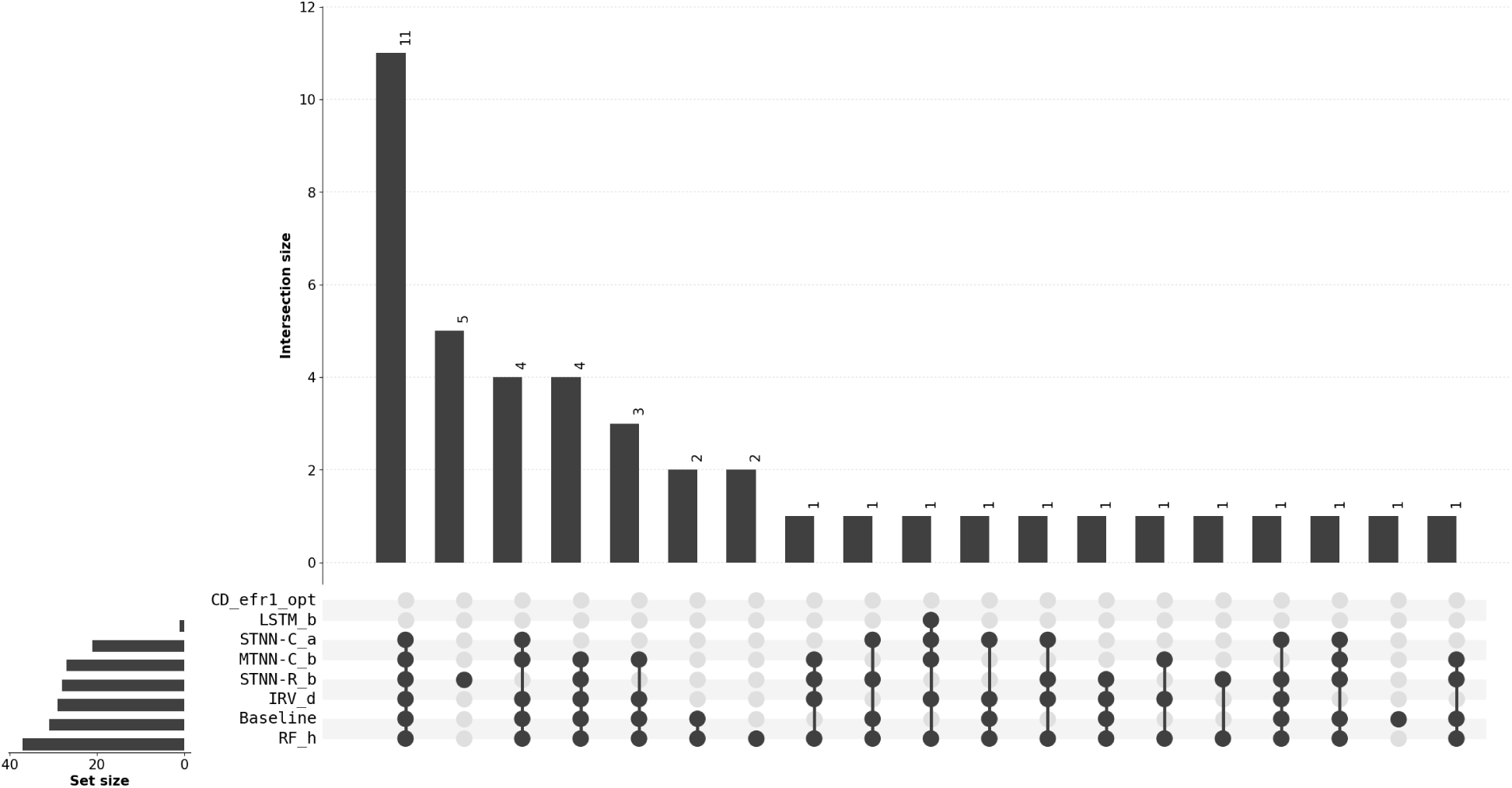
An UpSet plot showing the overlap between the top 250 predictions from the selected VS models and the chemical similarity baseline on **PriA-SSB prospective**. The plot generalizes a Venn diagram by indicating the overlapping sets with dots on the bottom and the size of the overlaps with the bar graph.^56^ Altogether, the combined predictions from the best-in-class VS methods and the baseline found 43 of the 54 actives.

The similarity baseline included one active compound that was excluded from the top 250 compounds from RF (Figure 4), but RF recovered a different member from this active compound’s SIM and MCS clusters (Appendix M).

Only the RF model recovered more active compounds in its top 250 predictions than the chemical similarity baseline including two unique actives not identified by any other model. Therefore, cross-validation with NEF_1%_ as the metric successfully identified the best PriA-SSB model before the prospective screen.

#### 3.2.1 Trained Models are Target-Specific

As a control, we retrained the best RF model on randomized data to confirm that its strong prospective performance was due to meaningful detected patterns among the active compounds instead of biases in the dataset. Similar to y-scrambling or y-randomization,^57^ we randomly permuted the binary activity labels in the **PriA-SSB AS** dataset, retrained the RF_h model on the randomized data, and evaluated the classifier on the **PriA-SSB prospective** dataset. This procedure was repeated 100 times with different y-scrambling performed each time. The number of active compounds in the top 250 predictions for these 100 runs are summarized in Figure S30. The mean number of actives was 0.83, and 55 of the runs found zero actives. The best y-scrambled run found only 10 actives, far less than the 37 actives when RF_h was trained on the real data.

In addition, we assessed the performance of all models trained on **RMI-FANCM FP** instead of **PriA-SSB AS** for making **PriA-SSB prospective** predictions on the new 22,434 compounds. As expected, the **RMI-FANCM FP** models perform poorly on **PriA-SSB prospective** (Table S29), indicating that the best **PriA-SSB AS** models have learned compound properties that are specific to PriA-SSB.

## 4 Discussion

We followed a VS pipeline with the goal of maximizing the number of active compounds identified in a prospective screen with a limited number of predictions. From an initial pool of structure-based and ligand-based models, we pruned models in a hyperparameter search stage and conducted cross-validation with multiple evaluation metrics. We used DTK+Means with the NEF_1%_ metric to select the best models based on the cross-validation results and experimentally evaluated their top 250 prospective predictions from a new library of 22,434 compounds. The single best model from our pool, which was RandomForest_h for **PriA-SSB AS**, was also the top performing model on **PriA-SSB prospective**. Therefore, our overall pipeline successfully identified the best prospective model.

Metrics like AUC[ROC] can compare models in general, regardless of cost or other additional constraints.^50^ However, for virtual screening in practice, one typically only experimentally tests a small fraction of all compounds in the test library. In this setting, metrics like EF that capture early enrichment are preferable. In our prospective screen, STNN-R_a had higher AUC[ROC] than RF_h (Appendix J), but the random forest found 8 more active compounds in its top 250 predictions (Appendix L). Our study suggests that EF*R*, or its normalized version NEF*R*, are the preferred metrics for identifying the best target-specific virtual screening method that maximizes *nhits* when there is a budget for experimental testing. Other metrics like AUC[ROC] or AUC[PR], which is more appropriate for problems where the inactive compounds far outnumber the actives,^58^ may still be reasonable for benchmarking virtual screening methods on large existing datasets where the entire ranked list of compounds is evaluated.^36^

Some recent studies^3,35,59^ reported that deep learning models substantially outperform traditional supervised learning approaches, including random forests. Our finding that a random forest model was the most accurate in both cross-validation and our prospective screen does not refute those results. Rather, it reinforces that the ideal virtual screening method can depend on the training data available, target attributes, and other factors. Therefore, careful target-specific cross-validation is important to optimize prospective performance. One cannot assume that deep learning models will be dominant for all targets and all virtual screening scenarios. We also recommend hyperparameter exploration for all models, including traditional supervised learning methods. For example, our best random forest model contained 8000 estimators, whereas a previous benchmark considered at most 50 estimators.^3^

Ramsundar et al.^14^ showed that performance improved in multi-task neural networks as they added more training compounds and tasks. Furthermore, the degree of improvement varied across the datasets and was moderately correlated with number of shared active compounds among the targets within a single dataset. Task-relatedness also affects the success of multi-task learning but is difficult to quantify.^60,61^ We observed that **PriA-SSB AS**, **PriA-SSB FP**, and **RMI-FANCM FP** have no shared actives with any of the PCBA tasks, and multi-task neural networks were not substantially better than single-task neural networks in **PriA-SSB AS** cross-validation (Figure 3). The MTNN-C model outperformed the STNN-C model in the prospective evaluation (Table 4), possibly because multi-task learning can help prevent overfitting,^62^ but was still considerably worse than the random forest. Multi-task random forests can also be constructed by using multi-task decision trees as the base learner.^63^ However, these methods have not been used widely in the context of virtual screening.

We focused on well-established machine learning models instead of more recent deep learning models, such as graph-based neural networks.^36,64–67^ This is because our main goal was to investigate the virtual screening principles for choosing the best model for a specific task (**PriA-SSB AS**) in a practical setting instead of broadly benchmarking virtual screening algorithms. In addition, a recent benchmark showed that conventional methods outperformed graph-based methods on most biophysics datasets.^36^

Consensus docking^48^ failed to recover any actives in the **PriA-SSB prospective** dataset, even though some of the individual docking programs did. Specifically for the PriA-SSB protein-protein interaction, docking is limited by the large, flat nature of the binding site. Many compounds that are inactive in the experimental screen have good scores and reasonable binding poses (per visual inspection) but fail to interrupt necessary specific interactions in the protein-protein interface. This will the limit overall performance by pushing true actives down the ranked list.

We use the docking results here only as comparison to common structure-based VS methods, though there are many other structure-based approaches and opportunities to train target-specific structure-based models. The individual docking and consensus methods do not make use of the initial HTS screening data, whereas the ligand-based machine learning methods do. A more direct comparison might be to re-train a custom consensus scoring function to include the initial HTS data, though this effort is out of scope for this study. In addition, there are computational tradeoffs between docking and ligand-based machine learning approaches. The machine learning models require substantial training time to select hyperparameters and fit models, but the trained models make predictions on new compounds very quickly. The docking programs take more time to score each new compound but have the advantage of not requiring training compounds.

The random forest model performed the best overall, but there were 6 active compounds identified by the other methods that the random forest missed (Figure 4). The single-task regression neural network recovered 5 of those 6 as well as unique active compound clusters (Appendix M). In addition, this regression model performed the best on **PriA-SSB FP** during cross-validation (Table 3), possibly because there are fewer binary actives in this dataset. In future work, we will explore whether ensembling classification and regression models, potentially in combination with structure-based VS algorithms, can further improve accuracy.

We emphasize our prospective performance on the new LC library, which minimizes the biases that make evaluation with retrospective benchmarks challenging.^68^ There are many sources of experimental error in HTS, and the active compounds in the prospective evaluation must still be interpreted conservatively. However, a VS algorithm that can prioritize compounds with high % inhibition in primary and retest screens is valuable for further compound optimization even if not all of the actives confirm experimentally. Our study provides guidelines for selecting a target-specific VS model and complements other practical recommendations for VS pertaining to hit identification, validation, and filtering^69^ as well as avoiding common pitfalls.^70^ Having established that our best virtual screening model successfully prioritized new active compounds in the LC library, another future direction will be to test prospective performance on much larger, more diverse chemical libraries.

## Acknowledgements

We acknowledge GPU hardware from NVIDIA; compute resources of the University of Wisconsin-Madison Center for High Throughput Computing; and funding from the Center for Predictive Computational Phenotyping NIH U54 AI117924, the University of Wisconsin Carbone Cancer Center Support Grant NIH P30 CA014520, and the Morgridge Institute for Research. Additional support for this research was provided by the University of Wisconsin-Madison Office of the Vice Chancellor for Research and Graduate Education with funding from the Wisconsin Alumni Research Foundation. We are grateful for the assistance and feedback from Chengpeng Wang, Haozhen Wu, and many members of the Center for High Throughput Computing and the Gitter lab. We thank Julio Lopes for alerting us about duplicate compounds in a preliminary version of the **PriA-SSB prospective** dataset.

## Supporting Information

Supporting Information available: Appendices A-N contain supporting text, Figures S1-S30, and Tables S1-S30.

